# Preserved Type 2 Immune Cell Plasticity in Human Obesity and Differential Immune Reconstitution After Bariatric Surgery

**DOI:** 10.64898/2026.05.09.723984

**Authors:** Jule Gawor, Andrea Deinzer, Maria Wick, Inaya Hayek, Christian Schwartz

**Affiliations:** Mikrobiologisches Institut - Klinische Mikrobiologie, Immunologie und Hygiene, Universitätsklinikum Erlangen and Friedrich-Alexander-Universität (FAU) Erlangen-Nürnberg, D-91054 Erlangen, Germany; FAU Immunomedicine (FAU I-MED), Erlangen, Germany

**Keywords:** obesity, type 2 immunity, bariatric surgery, eosinophils, macrophage polarization

## Abstract

**Background:** Obesity disrupts type 2 immune cell populations in white adipose tissue, replacing the homeostatic network of group 2 innate lymphoid cells (ILC2s), eosinophils, T helper 2 (Th2) cells, and alternatively activated macrophages (AAMs) with pro-inflammatory type 1 populations. Whether this remodelling reflects permanent immune impairment or a reversible shift in cellular equilibrium, and to what extent bariatric surgery restores type 2 immunity, remain incompletely understood.

**Methods:** We performed comprehensive immunophenotyping of visceral white adipose tissue (WAT) and peripheral blood from persons with severe obesity (people with obesity, PWO) scheduled for or having undergone bariatric surgery (sleeve gastrectomy, gastric bypass), combined with lean controls. Using flow cytometry, quantitative PCR, and *in vitro* polarization assays, we assessed immune cell frequencies, transcription factor expression, cytokine profiles, and functional polarization capacity across lean, pre-operative, and post-operative states.

**Results:** Obesity was associated with decreased eosinophil and CD8^+^ T cells frequencies in WAT, accompanied by an increase in CD4^+^ frequency and a shift from Th2 toward Th1 predominance, as well as elevated PD-1 expression on T cell subsets. Bariatric surgery partially normalised peripheral immune cell composition, reducing CD8^+^ T cell frequencies while increasing CD4^+^ T cells. Macrophage polarization capacity, dampened in pre-operative PWO, recovered after surgery. Conversely, Th2 polarization capacity and IL-13 production were reduced in post-operative T cells despite preserved function pre-operatively, indicating divergent trajectories of innate and adaptive immune reconstitution.

**Conclusion:** Type 2 immune cells retain functional plasticity in human obesity despite reduced frequency. Bariatric surgery differentially reconstitutes immune function, restoring macrophage plasticity while paradoxically reducing Th2 polarization capacity, arguing against uniform immune normalisation after weight loss.

**Funding:** German Federal Ministry of Research, Technology and Space (BMFTR, FKZ 01KI2109), Interdisciplinary Center for Clinical Research (IZKF, Faculty of Medicine, Friedrich-Alexander Universität (FAU) Erlangen-Nürnberg).

## Introduction

Obesity has reached epidemic proportions worldwide, with more than one billion individuals affected in 2022 [1]. The excessive accumulation of adipose tissue poses a major risk factor for many life-threatening comorbidities including type 2 diabetes mellitus, cardiovascular disease, certain types of cancer, arterial hypertension, and sleep apnoea [1, 2]. The health burden of obesity challenges healthcare systems globally, necessitating a deeper understanding of the underlying pathophysiological mechanisms that link excess adiposity to metabolic and systemic disease.

White adipose tissue (WAT) has been recognized not merely as an inert energy storage depot but as a dynamic endocrine and immunological organ [3]. WAT harbours various resident immune cell populations that play crucial roles in maintaining tissue homeostasis and metabolic health. These immune cells constitute a substantial fraction of the stromal vascular compartment and engage in complex bidirectional communication with adipocytes [4].

In lean healthy persons, adipose tissue maintains an anti-inflammatory microenvironment characterized by type 2 immune cell populations, including group 2 innate lymphoid cells (ILC2s), eosinophils, T helper 2 (Th2) cells, and alternatively activated macrophages (AAMs) [5-8]. These cells form an integrated network: ILC2s secrete IL-5 and IL-13 to recruit and sustain eosinophils, which produce IL-4 and IL-13 to polarize macrophages toward an alternatively activated phenotype [7, 8]. AAM support insulin sensitivity, while Th2 cells and regulatory T cells (Tregs) reinforce the anti-inflammatory milieu through additional IL-4, IL-13, and IL-10 secretion [9, 10]. The seminal study by Wu et al. demonstrated that eosinophils are the major IL-4-expressing cells in murine WAT and are required for sustaining AAMs associated with glucose homeostasis [8]. Subsequent work established ILC2s as critical upstream regulators of this circuit, with IL-33-driven ILC2 activation maintaining eosinophil recruitment and AAM polarization [11, 12].

Chronic overnutrition fundamentally disrupts this immune balance. Overweight and obesity trigger adipocyte hypertrophy, cellular stress, and hypoxia, initiating a cascade of pro-inflammatory events [13, 14]. Type 2 immune cell frequencies decline substantially while pro-inflammatory populations, including classically activated macrophages (CAMs), CD8^+^ T cells, and Th1 cells, accumulate in adipose tissue [15-18]. The landmark studies by Weisberg et al. and Xu et al. demonstrated that macrophages accumulate dramatically in obese adipose tissue, increasing from approx. 10% of the stromal-vascular cells in lean up to 50% in obese individuals, and represent the primary source of pro-inflammatory cytokines [13, 14]. Nishimura et al. subsequently showed that CD8^+^ effector T cell infiltration into obese WAT precedes macrophage accumulation, establishing T cells as early drivers of adipose inflammation [18].

This phenotypic shift from homeostatic type 2 toward pro-inflammatory type 1/3 immunity establishes chronic low-grade inflammation that contributes to insulin resistance and metabolic dysfunction [19, 20]. Importantly, this inflammatory remodelling extends beyond adipose tissue to affect immune cell populations systemically [21].

Within this framework, the PD-L1:PD-1 checkpoint has emerged as a critical regulator of adipose tissue immune homeostasis. We previously demonstrated that PD-L1 on dendritic cells limits T cell-mediated adipose tissue inflammation and ameliorates diet-induced obesity, establishing that immune checkpoints can actively maintain the type 2 environment in lean adipose tissue [22]. Whether PD-1 expression on T cell subsets in human adipose tissue inflammation reflects active immune regulation rather than exhaustion during obesity remains an important question.

Bariatric surgery, including sleeve gastrectomy and gastric bypass surgery, induces substantial and durable weight loss accompanied by remarkable metabolic improvements that often precede significant weight reduction [23]. Emerging evidence indicates profound immune system effects following surgery, including reduced T cell counts, decreased Th1/Th2 ratios, increased regulatory B cells, and improved NK cell activity [24-26]. However, critical questions remain: Does surgery fully restore the type 2 immune landscape of lean adipose tissue? Do systemic changes parallel local adipose remodelling? Recent murine studies by Cottam et al. reveal that adipose immune cells retain obesity-associated phenotypes despite weight loss with weight regain further exacerbating inflammatory signatures [27]. The concept of an “immunological scar” - persistent reprogramming of innate immunity triggered by past obesity - has been compellingly demonstrated by Hata et al. and Hinte et al., raising the question, whether human bariatric patients achieve complete immune reconstitution or exhibit persistent inflammatory imprints [28, 29].

To address these gaps, we analysed immune cell populations and functional responses in visceral adipose tissue and peripheral blood from individuals with obesity before and after bariatric surgery, and compared them to lean controls. We focussed on type 2 immune compartments - eosinophils, Th2 cells, and AAMs - testing the hypothesis that obesity depletes these populations without irreversibly impairing their functional capacity. Using flow cytometry, gene expression profiling, and in vitro polarization assays, we characterised both cell frequencies and functional potential. Our findings reveal that despite marked immune cell redistribution during obesity, type 2 immune cells retain robust functional responses to appropriate stimuli, while bariatric surgery induces partial but incomplete immune normalization with substantial inter-individual variation.

## Results

### Obesity remodels the immune cell landscape in visceral adipose tissue

To investigate the impact of obesity on resident immune cell populations, we performed flow cytometric analysis of the stromal-vascular fraction (SVF) from visceral WAT biopsies obtained during surgery from lean healthy controls (n=8) and people living with obesity (PWO; n=12). Clinical characteristics of our cohort are summarised in **Supplemental Table 1**.

Analysis of innate immune populations revealed that eosinophil (CD66b^+^Siglec-8^+^) frequencies were significantly (p<0.05) reduced in WAT of PWO compared to lean controls, consistent with published murine and human data demonstrating eosinophil depletion in obese adipose tissue (**Fig. 1A**) [8, 30]. Neutrophil (CD66b^+^Siglec-8^-^) and B cell (CD3^-^CD19^+^) frequencies showed a trend toward elevation in PWO but did not reach statistical significance. Monocyte (CD14^+^CD3^-^CD19^-^) frequencies were comparable between groups, with considerable interindividual variability.

**Figure 1.**
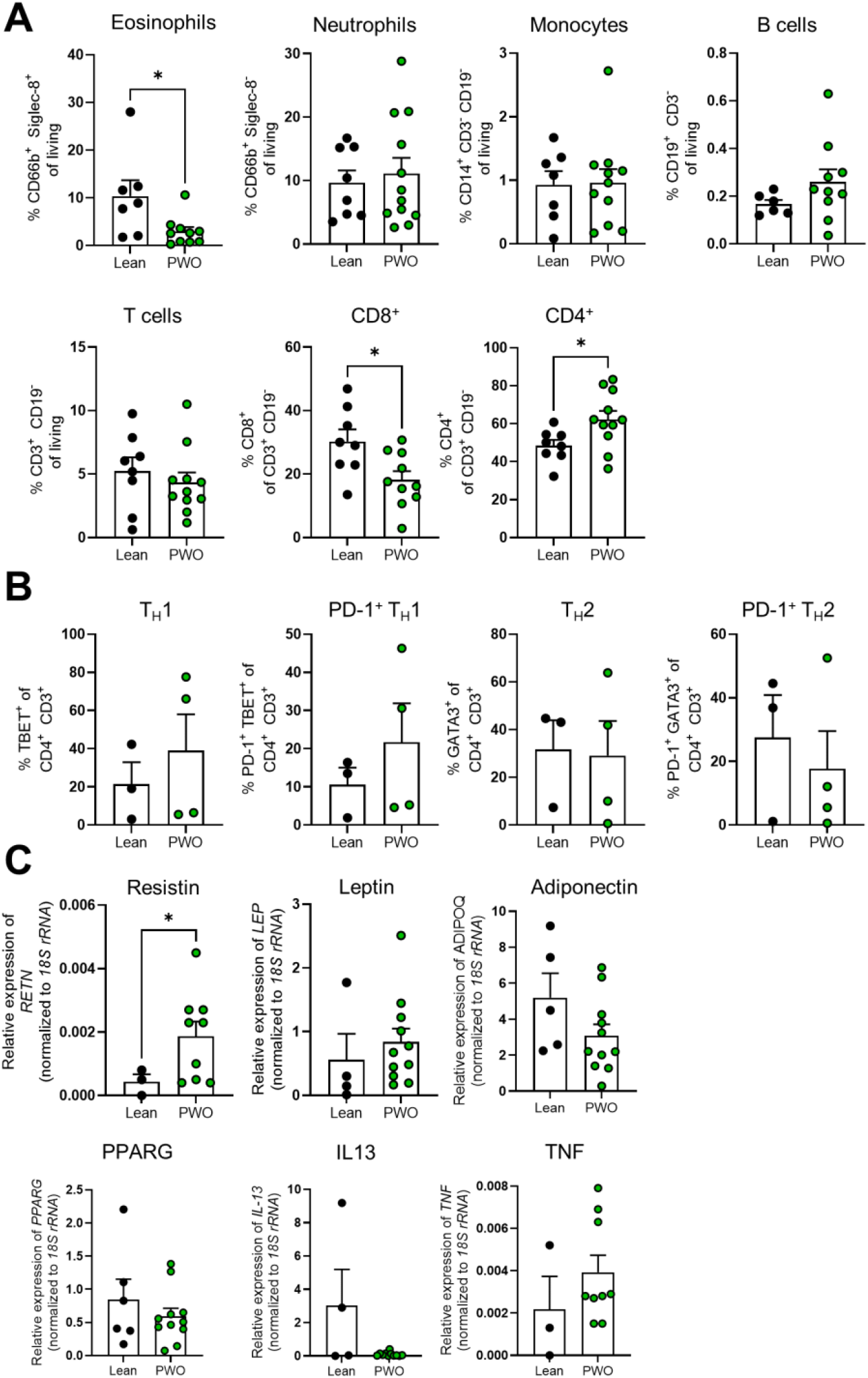
Obesity alters the immune cell composition within visceral adipose tissue. Visceral white adipose tissue (WAT) biopsies were obtained during surgery from lean healthy controls (n=8) and people living with obesity (PWO; n=12). **A, B)** Flow cytometric analysis was performed on the stromal-vascular fraction (SVF) of the WAT biopsies. **C)** Using qRT-PCR, the gene expression of Resistin (*RETN*), Leptin (*LEP*), Adiponectin (*ADIPOQ*), Peroxisome proliferator-activated receptor gamma (*PPARG*), Interleukin-13 (*IL13*) and Tumor Necrosis Factor (*TNF*) was analyzed. Relative expression was calculated using the 2^-ΔCt^ method, with human *18S rRNA* as a housekeeping gene. Mean ± SEM. Statistical outliers were identified and excluded using the ROUT method (Q=1%) in GraphPad Prism. *, p < 0.05; Mann-Whitney or Welch’s t test.

Within the adaptive immune compartment, CD8^+^ T cells were significantly decreased in WAT from PWO (p<0.05), while CD4^+^ T cells showed a significant increase relative to lean controls (p<0.05) (**Fig. 1A**). Detailed T cell subset analysis revealed a shift consistent with inflammatory polarization during obesity (**Fig. 1B**). Th1 cells, identified by TBET expression, were elevated in WAT from PWO. Notably, PD-1 expression on both Th1 and Th2 cells was detectable, with PD-1^+^ Th1 and PD-1^+^ Th2 populations showing trends toward differential regulation in obesity.

We further confirmed the metabolic and inflammatory remodelling associated with obesity by transcript analysis of whole WAT (**Fig. 1C**). Resistin and leptin expression were elevated in PWO, consistent with adipocyte hypertrophy and metabolic dysfunction. Adiponectin, a marker of healthy adipose function, trended lower in PWO. Peroxisome proliferator-activated receptor γ (PPARγ) expression, a key transcriptional regulator of adipogenesis and adipose tissue macrophage differentiation, showed a trend toward reduction in obesity. IL-13, a canonical type 2 cytokine, was reduced in PWO, while TNF expression was modestly elevated, indicating the shift from an anti-inflammatory to a pro-inflammatory tissue milieu.

### Circulating immune cell populations reflect systemic inflammation in obesity and partially normalise after bariatric surgery

We next examined whether immune alterations in obesity extend to circulating immune cells and assessed the impact of bariatric surgery on immune reconstitution. Flow cytometric analysis of peripheral blood from lean controls, pre-operative PWO, and post-operative PWO revealed distinct differences across groups (**Fig. 2A**).

**Figure 2.**
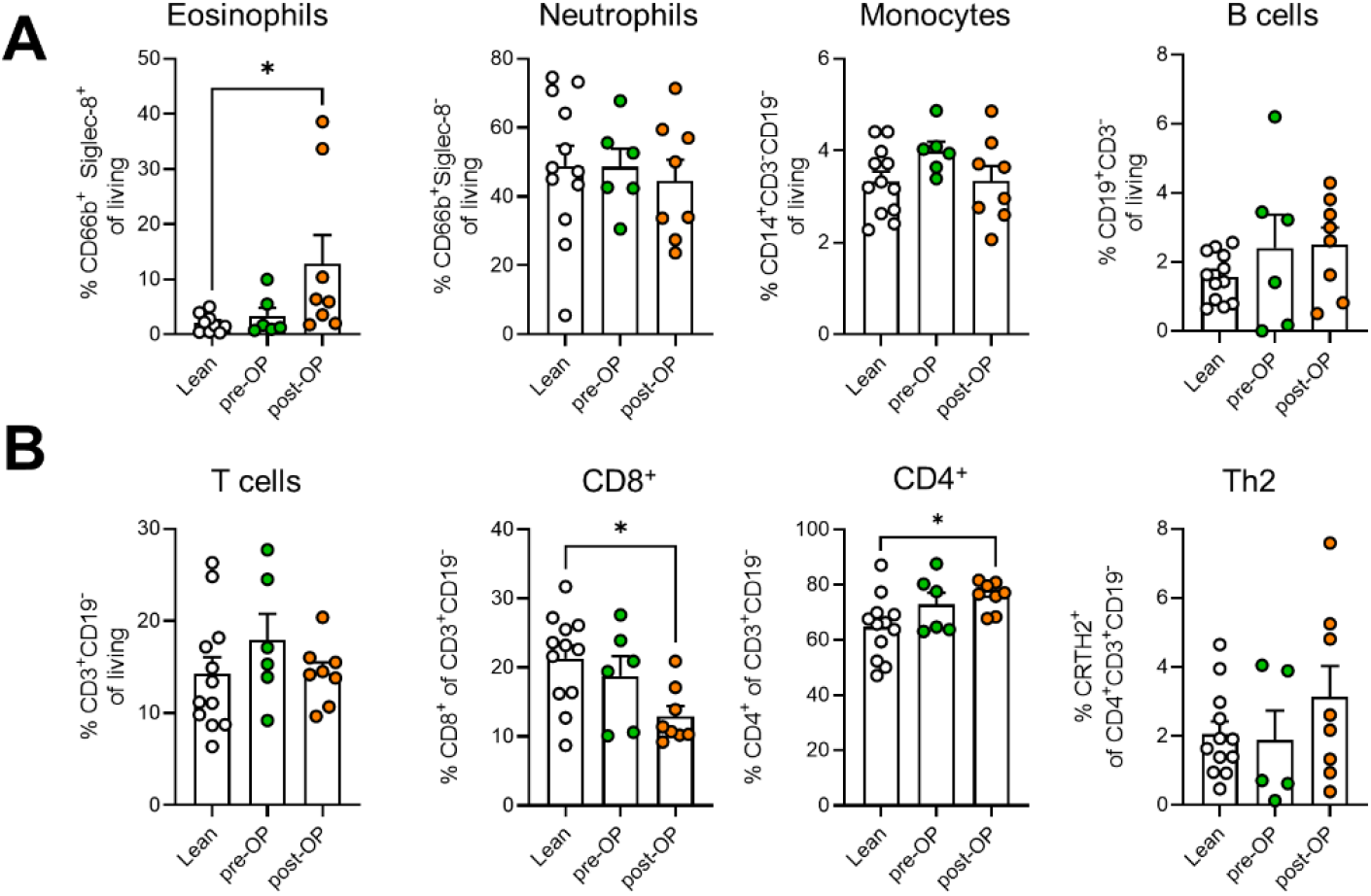
Circulating immune cell populations differ in obesity and normalize partially after bariatric surgery. **A, B)** Flow cytometric analysis of blood immune cells from anti-coagulated blood after depletion of erythrocytes with dextran and ACK lysis. Samples from lean, healthy donors (n=12) were compared to blood samples from PWO collected before (pre-OP; n=6) and 6 months after bariatric surgery (post-OP; n=8). Mean ± SEM. Statistical outliers were identified and excluded using the ROUT method (Q=1%) in GraphPad Prism. *, p < 0.05; Mann-Whitney or Welch’s t test.

Blood eosinophil frequencies showed variability across groups, with significant elevation in post-operative PWO compared to pre-operative PWO and lean controls. Neutrophil frequencies showed a high degree of interindividual variability. Monocytes showed a trend toward increased frequencies in pre-operative PWO, which returned to levels of lean controls post-surgery. Circulating B cell frequencies did not differ significantly between groups (**Fig. 2A**).

Within the T cell compartment, total T cell frequencies were comparable across groups, but important subset-specific changes emerged (**Fig. 2B**). Circulating CD8^+^ T cell frequency was comparable between lean controls and pre-operative PWO, but decreased significantly (p<0.05) following bariatric surgery. Conversely, CD4^+^ T cells increased after surgery. Interestingly, Th2 cells, identified by CRTH2 expression, showed a trend toward elevation post-surgery.

Paired analysis of pre- and post-operative blood samples (**Supplemental Figure 1**, n=2) showed that individual patients showed consistent decreases in neutrophil frequencies following surgery-induced weight loss. However, changes in eosinophils, T cells and B cells varied substantially between individuals, suggesting that immune reconstitution following bariatric surgery is heterogeneous and may depend on additional factors, such as degree of weight loss, metabolic improvement, and baseline immune status.

### T cells from PWO retain full capacity for Th2 polarization

Having observed altered T cell subset distribution in obesity, we investigated whether functional polarization capacity was affected. We isolated naive CD4^+^ T cells from the non-adherent cellular fraction of peripheral blood, activated them with anti-CD3/CD28 beads and IL-2, and cultured cells in Th1- or Th2-polarizing conditions for five days (**Fig. 3A**).

**Figure 3.**
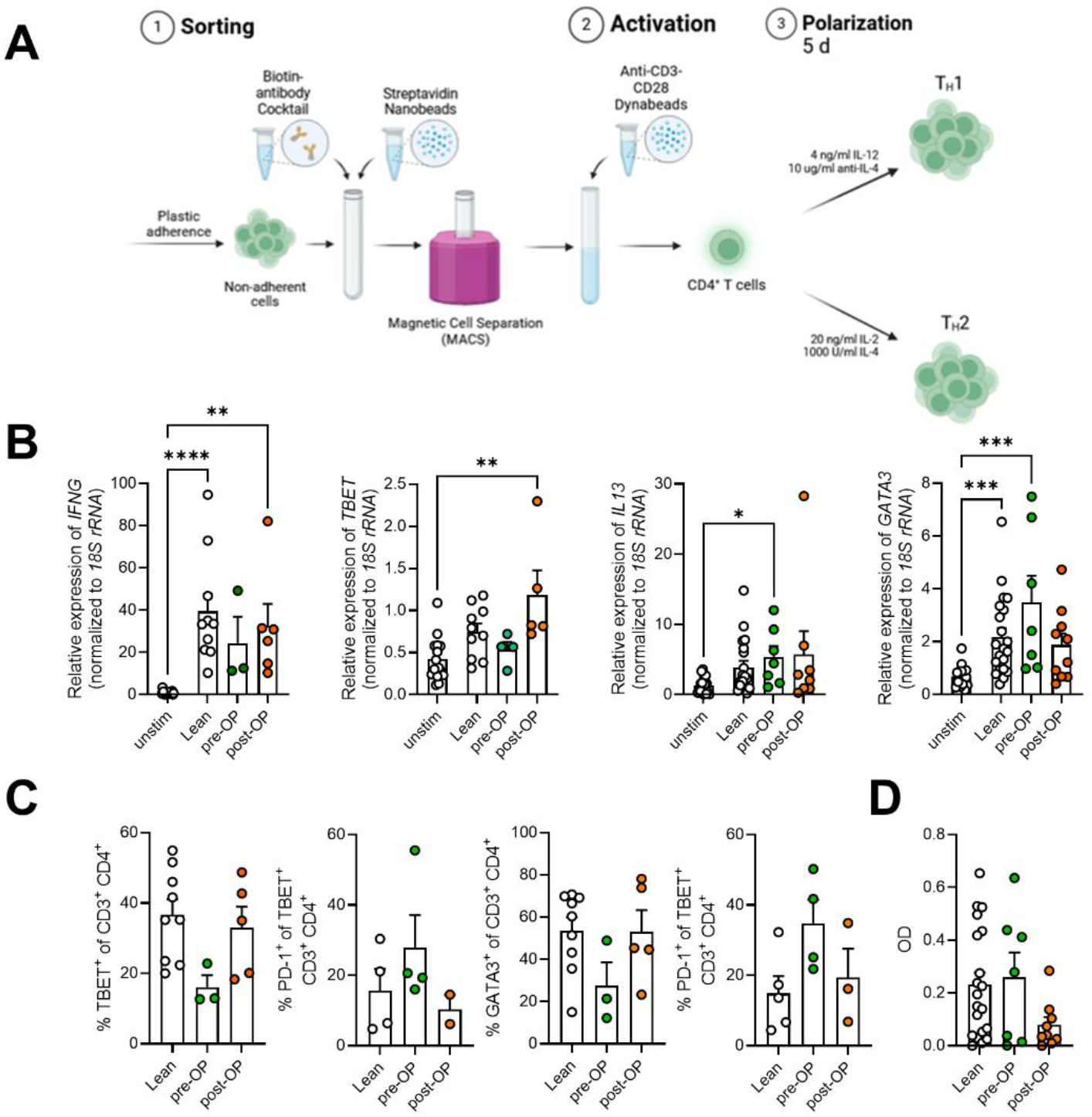
Th2 polarization potential is preserved in T cells derived from PWO. **A)** Schematic depiction of CD4^+^ T cell isolation and stimulation. Unstimulated cells cultured with IL-2 alone were included as a control. **B)** Expression of Interferon-γ (*IFNG*) and T-box transcription factor 21 (*TBET*) in Th1-polarized cells and IL-13 and GATA binding protein 3 (*GATA3*) in Th2-polarized cells. Graphs show the relative expression of genes (2^-ΔCt^) normalised to *18S rRNA*. Untreated cells from lean donors (n=15-20) were compared with polarized cells from lean donors (n=10-21), PWO before (pre-OP; n=3-7) and PWO after (post-OP; n=5-10) bariatric surgery. **C)** Flow cytometric analysis was performed from polarized cells. Relative protein expression from lean donors (n=4-9), pre-(n=3-4) and postoperative (n=2-5) patients are displayed. **D)** IL-13 secretion by Th2-polarized T cells was quantified by ELISA. Optical density (OD) was measured at 450 nm and compared between lean, pre- and post-operative samples (n=7-20). Mean ± SEM. Statistical outliers were identified and excluded using the ROUT method (Q=1%) in GraphPad Prism. ****, p < 0.0001; ***, p < 0.001; **, p < 0.01; *, p<0.05; One-Way ANOVA or Kruskal-Wallis-Test.

Gene expression analysis of polarized T cells revealed robust and equivalent functional responses across groups (**Fig. 3B**). Th1 polarization induced marked upregulation of IFN-γ mRNA in cells isolated from lean, pre-operative and post-operative donors, with no significant difference between lean and PWO groups. TBET expression similarly increased following Th1 polarization, confirming intact Th1 programming. Critically, Th2 polarization was equally preserved. IL-13 mRNA was significantly upregulated upon IL-4 stimulation with comparable induction in lean, pre-operative, and post-operative PWO cells. GATA3, the master transcription factor for Th2 differentiation, was significantly induced by IL-4 (p<0.05), with no impairment in PWO-derived T cells. Post-operative samples showed a trend towards reduction in Th2 polarization, indicating that surgical weight loss may reduce this functional property.

Flow cytometric validation confirmed the gene expression findings (**Fig. 3C**). TBET and GATA3 protein expression among CD4^+^ T cells reflected the expected polarization patterns, with no deficit in pre-OP PWO-derived cultures and reduced GATA3-expression in post-operative T cells. Interestingly, PD-1 co-expression with TBET returned to lean levels in T cells derived from post-OP donors, while in GATA3^+^ T cells expression remained elevated. In order to assess functionality of polarized T cell subsets, we measured cytokine production by ELISA (**Fig. 3D**). IL-13 secretion from Th2-polarized cells was comparable between lean controls and pre-OP PWO, demonstrating that the type 2 cytokine production machinery remains intact in T cells from PWO. Importantly, after surgery, we consistently observed reduced GATA3 expression and limited production of IL-13 after Th2 polarization.

### Eosinophils from PWO maintain robust IL-5 responsiveness

Eosinophils play critical roles in type 2 immunity and adipose tissue homeostasis [8, 30, 31]. We isolated blood eosinophils from lean controls and PWO and characterized their responses to IL-5, a key eosinophil survival and activation factor (**Fig. 4**).

**Figure 4.**
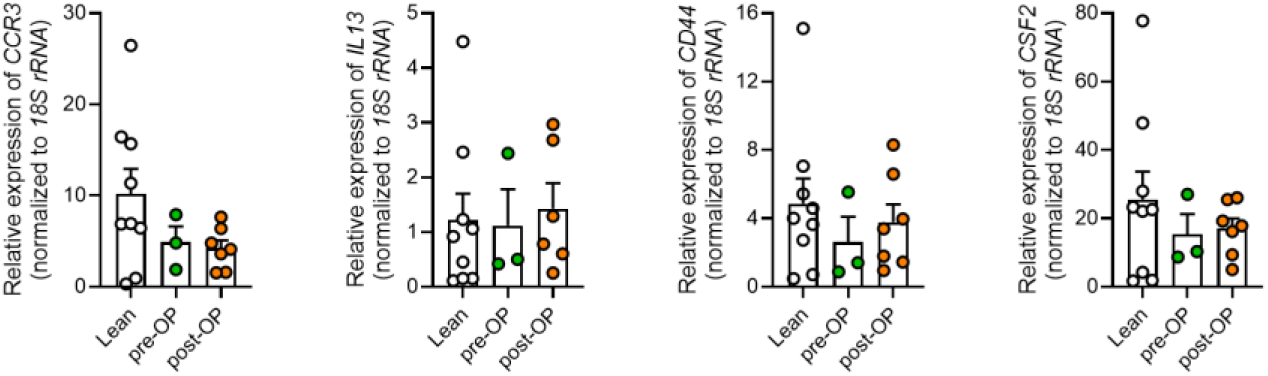
Eosinophil function and responsiveness to IL-5 are not impaired during obesity. Eosinophils were isolated from peripheral blood and stimulated with IL-5 for 24 hours. Transcript expression of C-C chemokine receptor type 3 (*CCR3*), *IL13*, Cluster of differentiation 44 (*CD44*) and Colony stimulating factor 2 (*CSF2*). Expression levels were compared between lean donors (n = 9) and PWO before (pre-OP; n = 3) and after (post-OP; n = 7) bariatric surgery. Mean ± SEM. Statistical outliers were identified and excluded using the ROUT method (Q=1%) in GraphPad Prism. One-Way ANOVA or Kruskal-Wallis-Test.

Gene expression analysis demonstrated that eosinophils from PWO maintained robust responses to IL-5 stimulation. CCR3, a chemokine receptor mediating eosinophil tissue recruitment, was expressed across all groups with lower expression in IL-5 stimulated cells from PWO. IL-13 showed variable expression levels with maintained responsiveness in PWO. CD44, an activation marker associated with tissue-resident eosinophils, and CSF2 (GM-CSF), which is expressed mainly by inflammatory eosinophils, were expressed across all groups with no significant impairment in PWO (**Fig. 4**).

These findings indicate that despite systemic inflammation in obesity, the core eosinophil functional programs, including chemokine receptor expression, type 2 cytokine production, and survival signalling, remain intact and responsive to IL-5-induced activation ex vivo.

### Monocyte-derived macrophages from PWO retain AAM polarization capacity with persistent CAM

*bias* Adipose tissue macrophages are key mediators of obesity-induced inflammation. To determine whether circulating monocytes from PWO have altered polarization capacity, we generated monocyte-derived macrophages in vitro and exposed them to CAM-polarizing (LPS+IFN-γ) or AAM-polarizing (IL-4) stimuli (**Fig. 5A**).

**Figure 5.**
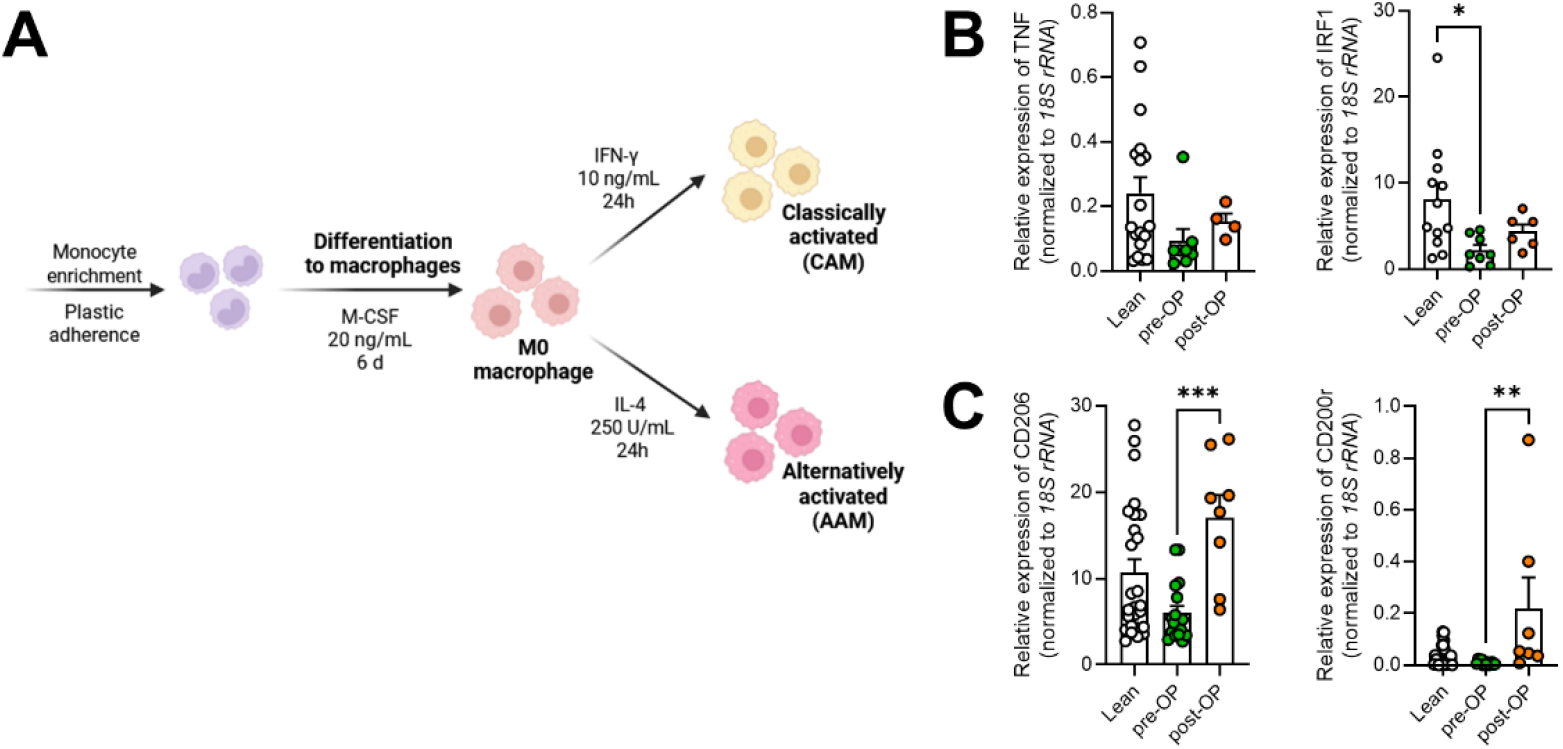
Monocyte-derived macrophages regain polarization capacity after bariatric surgery. **A)** Schematic depiction of monocyte (isolated from peripheral blood) differentiation into macrophages and their subsequent polarization into classically activated macrophages (CAM) or alternatively activated macrophages (AAM). **B)** Gene expression in CAM-polarized cells was quantified by qRT-PCR and compared among lean donors (n = 12-18), preoperative (n = 7-8), and postoperative samples (n = 4-6) from PWO. Relative expression of TNF and interferon regulatory factor 1 (IRF1) was normalised to *18S rRNA*. **C)** Gene expression in AAM-polarized cells was quantified by qRT-PCR and compared among lean donors (n = 23-26), preoperative (n = 13-18), and postoperative samples (n=8) from PWO. Relative expression of mannose receptor C-type 1 (*CD206*) and CD200 receptor 1 (*CD200R1*) was normalised to *18S rRNA*. Mean ± SEM. Statistical outliers were identified and excluded using the ROUT method (Q=1%) in GraphPad Prism. ***, p < 0.001; **, p < 0.01; *, p<0.05; One-Way ANOVA or Kruskal-Wallis-Test.

Unexpectedly, analysis of the CAM-associated marker TNF revealed a trend toward elevated expression in lean macrophages (**Fig. 5B**). IFN-γ-stimulation led to increased TNF transcript levels in macrophages from lean controls compared to pre-operative PWO. After surgery, macrophages regained responsiveness toward IFN-γ stimulation. Similarly, IRF1, a key IFN-γ signalling mediator, showed regained upregulation following bariatric surgery.

Analysis of the canonical AAM marker CD206 (mannose receptor) demonstrated that IL-4 induced substantial upregulation in macrophages from lean controls (**Fig. 5C**). Macrophages derived from pre-OP PWO showed decreased capacity to respond to IL-4, with lower expression of CD206 and CD200R. However, following bariatric surgery both AAM markers were significantly increased compared to pre-operative samples.

Collectively, these results demonstrate that macrophages derived from pre-operative PWO show decreased polarization capacity both toward CAM and AAM subsets. Importantly, post-bariatric surgery macrophages regained responsiveness towards cytokine stimulation.

## Discussion

This study provides a comprehensive analysis of type 2 immune cell frequencies and functional capacity in human obesity and following bariatric surgery. Our key finding is that despite marked alterations in type 2 immune cell populations in visceral adipose tissue and peripheral blood during obesity, the core functional machinery of Th2 cells and eosinophils remains intact, while macrophages become unresponsive to cytokine mediated polarization in vitro. Bariatric surgery partially normalises immune cell composition but does not fully restore the lean immune phenotype within the timeframe examined. These observations have important implications for understanding obesity-associated immune dysfunction and for developing targeted immunomodulatory therapies.

### Type 2 immune circuit disruption in human adipose tissue

Our observation of reduced eosinophil frequencies in obese WAT aligns with the growing body of evidence establishing eosinophils as critical regulators of adipose tissue homeostasis. Wu et al. demonstrated that eosinophils sustain AAMs in adipose tissue and that their absence exacerbates metabolic dysfunction [8]. More recently, Hernandez et al. reported significantly reduced adipose tissue eosinophil content in humans with obesity, with concurrent decreases in IL-4 expression and eotaxin family members [30]. Our data extend these observations by demonstrating that eosinophil depletion occurs alongside a broader restructuring of the WAT immune landscape, including decreased CD8^+^ T cells and a shift from Th2 toward Th1 predominance.

The concurrent decrease of CD8^+^ T cells in obese WAT contrasts with the seminal finding by Nishimura et al. that CD8^+^ effector T cell infiltration precedes macrophage accumulation in murine adipose tissue [18]. Our data indicate that in established human obesity, CD8^+^ T cell numbers decline while CD4^+^ T cells accumulate - a pattern that may reflect species-specific differences, the chronicity of human obesity compared to short-term murine high-fat diet models, or T cell exhaustion and apoptosis in the chronically inflamed tissue milieu. Porsche et al. recently described adipose tissue T cell exhaustion in human obesity [32], which could account for the decreased CD8^+^ compartment we observe. The elevated PD-1 expression on both Th1 and Th2 cells in our WAT data connects to our prior work demonstrating that the PD-L1/PD-1 axis on dendritic cells restrains T cell-mediated adipose inflammation [22], and supports a model in which checkpoint engagement is actively attempting to limit ongoing inflammatory damage.

### Preserved Th2 and eosinophil functional capacity as a therapeutic opportunity

The central translational finding of our study is that Th2 cells and eosinophils from individuals with obesity retain robust functional capacity when provided with appropriate polarizing signals *in vitro*. This challenges a deterministic view of obesity-associated immune dysfunction and supports a model in which the local microenvironment, rather than irreversible cellular reprogramming, is the primary driver of impaired type 2 responses.

For T cells, our demonstration that naive CD4^+^ cells from pre-operative PWO undergo normal Th2 polarization with equivalent IL-13 and GATA3 induction is notable. The metabolic milieu in obese adipose tissue is characterised by high leptin, low adiponectin, elevated free fatty acids, and IFN-γ dominance and may suppress Th2 differentiation through mTORC1 activation and inhibition of GATA3 transcription [20, 33]. Winer et al. demonstrated that adoptive transfer of CD4^+^ T cells with intact Th2 capacity can reverse insulin resistance in obese mice [10]. Our data suggest that this approach may be feasible in humans, as the cellular machinery for Th2 responses remains intact in established obesity. Intriguingly, post-operative T cells showed reduced Th2 polarization capacity with decreased GATA3 expression and IL-13 production, suggesting that surgical weight loss and the associated metabolic shifts may paradoxically diminish type 2 immune polarization capacity, which warrants further investigation. For eosinophils, maintained IL-5 responsiveness with preserved CCR3, IL-13, and CSF2 expression indicates that the eosinophil activation programme remains functional. Calco et al. have emphasised that eosinophil function, rather than number alone, is the critical determinant of metabolic outcomes [31].

Thus, therapeutic strategies to increase eosinophil numbers in the WAT of PWO may merit further investigation as we now show that ex vivo eosinophils retain their function.

### Macrophage hyporesponsiveness in obesity and recovery after bariatric surgery

In contrast to the preserved functional capacity of Th2 cells and eosinophils, monocyte-derived macrophages from pre-operative PWO showed a striking reduction in polarization capacity toward both CAM and AAM phenotypes. This functional dampening - with reduced TNF induction upon IFN-γ stimulation and impaired CD206 and CD200R upregulation upon IL-4 exposure - suggests a state of macrophage tolerance or exhaustion rather than the enhanced inflammatory priming classically attributed to obesity. This finding is unexpected in light of the trained immunity framework, in which metabolic signals such as oxidised LDL and saturated fatty acids are thought to enhance innate immune responsiveness through epigenetic reprogramming [34, 35]. One possible explanation is that the chronic exposure of monocytes to inflammatory mediators in established obesity induces a tolerogenic state analogous to endotoxin tolerance, in which prolonged stimulation leads to functional hyporesponsiveness rather than priming [36, 37]. Consistent with this interpretation, a recent study demonstrated that palmitic acid and conditioned medium from obese adipose tissue induce TLR4-dependent trained immunity in bone marrow-derived macrophages, an effect that was abolished by methyltransferase inhibition [38]. The discrepancy between trained immunity induced by acute lipid exposure and the functional dampening we observe in monocyte-derived macrophages from PWO with long-standing obesity suggests that chronic metabolic stress may push macrophages past the priming window into a tolerogenic state. Alternatively, the in vitro differentiation protocol, which removes monocytes from their inflammatory milieu for six days of M-CSF-driven maturation, may unmask an intrinsic functional deficit that is partially compensated in vivo by the abundant inflammatory signals present in obese tissue.

Importantly, macrophages derived from post-operative PWO regained cytokine responsiveness, with restored TNF induction upon IFN-γ stimulation and improved CD206 upregulation upon IL-4 exposure. This recovery of macrophage plasticity following weight loss suggests that the obesity-associated dampening of macrophage function is at least partially reversible and may be driven by circulating factors, such as elevated leptin, insulin, or free fatty acids, that resolve following bariatric surgery. The differential trajectory of macrophages compared to T cells after surgery (macrophage recovery versus reduced Th2 capacity) highlights the distinct mechanisms governing innate versus adaptive immune reconstitution and argues against a uniform model of post-surgical immune normalisation.

### Incomplete immune reconstitution following bariatric surgery

Bariatric surgery partially normalised peripheral immune cell composition in our cohort, with notable changes in CD8^+^ and CD4^+^ T cell frequencies. These findings are consistent with CyTOF analysis of morbidly obese following surgery, who reported that naïve T cells did not fully recover within 9-11 months after surgery [24], and with Wijngaarden et al., who found that T cell differentiation profiles and cytokine-producing capacity remained altered despite B cell normalisation [26]. The substantial interindividual variability in paired analyses underscores that immune reconstitution following bariatric surgery is heterogeneous and likely depends on the degree of weight loss, metabolic improvement, duration of prior obesity, and baseline immune status.

The concept of an “immunological scar” from obesity has gained support from recent studies. Hata et al. demonstrated that diet-induced obesity triggers persistent epigenetic reprogramming of the innate immune system through stearic acid and TLR4 signalling [28], and Hinte et al. showed that human adipose tissue retains transcriptional and epigenetic alterations even years after bariatric surgery [29]. Most recently, the adipose niche at single-nucleus resolution across lean, obese, and weight-loss states was mapped and the study found that senescence in adipocyte progenitors and vascular cells is potently reversed by weight loss, whereas other cellular and molecular alterations persist [39]. Cottam et al. used CITE-seq to demonstrate that obesity-induced imprinting of adipose immune cells persists through weight loss, with impaired recovery of Tregs and ILC2s [27]. Our data add a nuanced perspective: while the immunological scar may alter macrophage responsiveness and tissue immune cell composition, the intrinsic Th2 and eosinophil differentiation potential appears resistant to permanent reprogramming. This distinction has therapeutic implications: interventions that shift the local microenvironment toward type 2-favouring conditions may be sufficient to re-engage the preserved functional capacity of immune cells without requiring reversal of epigenetic changes.

## Limitations

Several limitations warrant consideration. Lean WAT controls were obtained from patients undergoing elective abdominal surgery rather than healthy volunteers, representing the standard approach given that obtaining WAT from truly healthy individuals is ethically not feasible [30]. The age difference between lean controls (mean 30.6 years) and PWO (mean 48.0 years) is a potential confounder, though age-related immunosenescence would bias against our key finding of preserved Th2 capacity [40]. Paired pre- and post-operative analyses were limited to a subset of patients; prospective studies with extended follow-up are needed to capture the full kinetics of immune reconstitution. In vitro polarization assays assess maximum differentiation capacity under supraphysiological conditions and may not fully recapitulate the in vivo milieu.

## Conclusions

Our data support a model in which obesity creates a quantitative deficit in type 2 immune cells and induces macrophage hyporesponsiveness without abolishing the intrinsic functional potential of Th2 cells and eosinophils. Bariatric surgery differentially restores immune function through recovering macrophage plasticity, while paradoxically reducing Th2 capacity. Strategies to boost type 2 immunity, such as IL-33 administration, eotaxin pathway modulation, or GLP-1 receptor agonists with emerging immunomodulatory properties [41], represent promising avenues for complementing the metabolic benefits of bariatric surgery. The preserved functional plasticity of type 2 immune cells provides a rational foundation for such approaches.

## Methods

### Study design and participants

#### Ethics declaration

This study was conducted in accordance with the Declaration of Helsinki and approved by the local ethics committee. All participants provided written informed consent prior to enrolment. Participation was voluntary and without financial compensation.

#### Patient cohorts

In collaboration with the Adipositaszentrum Erlangen, we enrolled 33 patients from the clinic and 29 control individuals following provision of written informed consent. Twenty-five patients were diagnosed with severe obesity (BMI ≥35 kg/m^2^, grade II, or ≥40 kg/m^2^, grade III) and were scheduled for bariatric surgery or had previously undergone bariatric surgery. These individuals are referred to as people with obesity (PWO). Surgical procedures included sleeve gastrectomy (n=14), gastric bypass (n=8), sleeve gastrectomy revision (n=2), and intragastric balloon explantation (n=1).

The mean age of the PWO cohort was 48.0 ± 8.9 years. Mean weight before surgery was 171.32 ± 34.6 kg (BMI 54.3 kg/m^2^) and decreased to 132.3 ± 22.5 kg (BMI 43.6 kg/m^2^) after surgery, corresponding to a mean weight reduction of 32.00 ± 14.1 kg (-10.2 kg/m^2^ BMI). The cohort exhibited expected obesity-related comorbidities, including arterial hypertension (60.0%), exertional dyspnoea or obesity-hypoventilation syndrome (32.0%), orthopaedic disorders (68.0%), and metabolic disorders, such as type 2 diabetes (40.0%) (**Supplemental Table 1**).

For the control group, blood samples were obtained from 14 male and 15 female individuals aged 22-61 years (mean 30.6 ± 5.9 years) with a BMI <30 kg/m^2^ (mean 23.5 ± 2.6 kg/m^2^). Non-obese controls had no documented metabolic disorders or chronic inflammatory conditions.

For each donor, we aimed to perform the full panel of analyses; however, limited sample volumes, particularly for adipose tissue biopsies and pre-operative blood draws, precluded all assays in every individual, resulting in variable sample sizes across experiments.

### Sample collection and preparation

#### Visceral adipose tissue

Visceral adipose tissue was obtained during laparoscopic surgery, stored in Ringer’s solution at 4°C, and processed within 24 hours. A total of 12 samples were collected from PWO, and 8 samples were obtained from lean individuals who underwent either reflux surgery (n=6) or gastric pacemaker implantation (n=2). The lean cohort had a mean age of 57.0 ± 20.0 years (range 27-86 years) and a mean BMI of 24.33 ± 3.1 kg/m^2^.

Adipose tissue was weighed and divided using sterile scissors for downstream analyses. 0.1-0.2 g of tissue was snap-frozen in liquid nitrogen and stored at -80°C for subsequent gene expression analysis via quantitative PCR. An equal amount was transferred into a 50 ml Falcon tube containing 5 ml of complete RPMI medium. The tissue was mechanically minced with scissors, and 100 µl of Collagenase D (50 mg/ml; final concentration: 1 mg/ml) was added for enzymatic digestion. Samples were incubated at 37°C for 45 min with shaking at 200 rpm. Following digestion, the cell suspension was passed through a 100 µm cell strainer, centrifuged, and red blood cells were lysed using ammonium-chloride-potassium (ACK) lysis. The remaining cells were counted, resuspended at a concentration of 0.5 × 10^6^ cells per 100 µl, and transferred to a FACS staining plate for subsequent staining.

#### Peripheral blood samples

Blood samples were collected from PWO at two time points: the initial blood sample was collected within one year prior to surgery and designated the pre-operative sample (median 43 days before surgery; range 2-344 days). The post-bariatric sample was collected between six months and 18 months after surgery (4 samples >18 months), designated the post-operative sample (median 334 days after surgery; range 220-2450 days). At each time point, 23.5 ml of EDTA-anticoagulated blood (S-Monovetten Kalium-EDTA, Sarstedt) was collected and processed within 24 hours. For the control group, 20 ml citrate-anticoagulated blood (S-Monovette Citrat 9NC, Sarstedt) was collected and processed within 24 hours of collection.

#### Whole blood immune cell isolation

In order to analyse blood immune cell distribution, 2 ml of anticoagulated blood was transferred to a 15 ml centrifuge tube (Corning or Sarstedt) and centrifuged (5 min, 400 × g, room temperature [RT]). After removal of plasma, the remaining pellet was resuspended in Dulbecco’s phosphate-buffered saline (DPBS; DPBS w/o Ca^2+^ and Mg^2+^, PAN Biotech) to a final volume of 5 ml. Subsequently, the cell-DPBS mixture was mixed with an equal volume of 3% dextran solution (Dextran 500, Roth) and incubated at RT for 30 minutes to sediment erythrocytes. Following two rounds of ACK lysis, cells were resuspended in 200 µl RPMI medium (Roswell Park Memorial Institute [RPMI] 1640 Medium, Gibco/Life Technologies) for subsequent flow cytometric staining.

#### Isolation of peripheral blood mononuclear cells

Peripheral blood mononuclear cells (PBMCs) were isolated from 15 ml of anticoagulated blood using density gradient centrifugation. After incubation with 3% dextran solution, the erythrocyte-depleted supernatant was diluted with DPBS to a total volume of 35 ml and carefully layered on top of 15 ml Biocoll (Bio&Sell, Feucht, Germany) in a fresh 50 ml tube. Density gradient centrifugation was performed at 1,000 × g for 15 min at RT without brake. After centrifugation, the plasma layer was discarded, and the thin white interphase containing leukocytes was carefully collected and washed twice with PBS to improve purity. Cells were counted and resuspended at 1 × 10^7^ cells per 500 µl in human dendritic cell (hDC) medium (RPMI 1640 supplemented with 1% HEPES (pH 7.3), 1% human serum albumin (hSA), 1% L-glutamine, 1% penicilllin-streptomycin) for monocyte and T cell isolation, or cryopreservation. For cryopreservation, cells were centrifuged and resuspended in 500 µl 20% human serum albumin (hSA) in a cryo-tube, then incubated for 10 min on ice. Subsequently, 500 µl of freezing medium (55% hSA, 25% Glucose 40%, 20% dimethyl sulfoxide (DMSO)) was added, the tubes were wrapped in paper, and stored at −80°C.

#### Monocyte-derived macrophages

Monocytes were isolated based on their adherent behaviour. A total of 500 µl of the PBMC suspension was mixed with 1,500 µl hDC medium in 6-well plates (Thermo Scientific) and incubated for 75 min at 37°C and 5% CO_2_. Afterwards, non-adherent cells were removed and placed on ice for subsequent T cell isolation. Each well was washed three times with 2 ml RPMI. To induce differentiation into macrophages, adherent monocytes were cultured in 2 ml human macrophage medium (RPMI 1640 supplemented with 1% HEPES (pH 7.3), 10% hSA, 1% L-glutamine, 1% penicillin-streptomycin) supplemented with 20 ng/ml macrophage colony-stimulating factor (M-CSF) for 6 days (37°C, 5% CO_2_). On day 3, 500 µl of fresh M-CSF-containing medium was added to each well.

Following the 6-day incubation period, during which monocytes differentiated into macrophages, cells were detached from the plate, washed, counted and seeded into 96-well flat-bottom plates (Thermo Scientific) at a concentration of 2×10^6^/ml. Cells were either left unstimulated (control), or stimulated with interferon-γ (IFN-γ, 10 ng/ml) or interleukin-4 (IL-4, 250 U/ml). After 24h, supernatants were collected and stored at -20°C, and cells were harvested for gene expression analysis and flow cytometry.

#### T cell isolation and polarization

CD4^+^ T cells were magnetically isolated from the non-adherent cellular fraction using the Mojosort Human CD4 Naïve T Cell Isolation Kit (BioLegend) according to the manufacturer’s protocol. 2 × 10^5^ naïve T helper cells (>95% purity) were activated with 4 × 10^6^ Dynabeads Human T-Activator (Thermo Scientific) and 40 ng/ml recombinant human IL-2 (ImmunoTools, 20 µg/ml). Cells were either left unstimulated or supplemented with 20,000 U/mL IL-4 (Miltenyi Biotec, 2×10^5U/ml) for Th2 polarization, or 8 ng/ml IL-12 (Peprotech, Fisher Scientific, 10 µg/ml) and 10 µg/ml anti-IL-4 (BioCell, 5,72 mg/ml) antibodies for Th1 polarization. On day 3, half the medium was replaced with fresh IL-2-containing hDC medium. After 5 days of incubation (37°C, 5% CO_2_), supernatants were collected and stored at −20°C, and cells were harvested for gene expression analysis and flow cytometry staining.

#### Eosinophil isolation

To isolate eosinophils, the EasySep Direct Human Eosinophil Isolation Kit (StemCell Technologies) was used according to the manufacturer’s protocol. The purified cells were counted and resuspended in eosinophil culture medium (RPMI 1640 supplemented with 20% fetal calf serum (FCS), 25 mM HEPES, 1% penicillin-streptomycin, 2 mM L-glutamine, 10 mM ß-mercaptoethanol, 10 mM non-essential amino acids). Isolation purity was assessed by flow cytometry (purity >95%). 10^5^ eosinophils were seeded per well and stimulated with either 20 ng/ml IL-5 (R&D Systems) or left untreated. After 24 h (37°C, 5% CO_2_), eosinophils were harvested in 300 µl RNAprotect Cell Reagent (Qiagen) for downstream RNA analysis. For histological analysis, eosinophils were cultured in chamber slides using the same culture conditions.

#### Flow cytometry

For flow cytometry analysis, 10^6^ cells were stained with Zombie aqua fixable dye (BioLegend) in PBS for 30 minutes at 4°C. Cells were washed with PBS buffer. Surface staining was performed using antibodies listed in **Supplemental Table 2** diluted in FACS buffer (PBS, 2% FCS (Sigma Aldrich) and NaN_3_) and incubated for at least 30 minutes at 4°C.

For subsequent transcription factor staining in T cells, surface-stained cells were fixed and permeabilized using FoxP3 staining buffer kit (Thermo Scientific). After incubation for 30 min at 4°C, cells were washed with Perm/Wash buffer diluted 1:10 in PBS. Cells were stained overnight (4°C) with antibodies listed in **Supplemental Table 2** diluted in Perm/wash buffer. Cells were washed, filtered through a 100 µm nylon mesh (Franz Eckert GmbH), and acquired on a BD LSRFortessa flow cytometer.

#### Gene expression analysis

RNA was isolated from eosinophils using the RNeasy Mini kit (Qiagen), or from T cells and macrophages using phenol/chloroform extraction with TRIzol Reagent (Thermo Scientific). Concentration and purity were measured using a NanoDrop spectrophotometer (Thermo Scientific). For each sample, a total of 500 ng RNA was reverse transcribed into cDNA using the High-Capacity Reverse Transcription Kit (Thermo Scientific) according to the manufacturer’s instructions. Real-time quantitative PCR was performed on a ViiA7 Real-Time PCR System (Applied Biosystems) using the SYBR Select Mastermix kit (Applied Biosystems) according to the manufacturer’s protocol. Primer sequences are listed in **Supplemental Table 3**. 18S ribosomal RNA (*18S rRNA*) or ribosomal protein 13A (*RPL13A*) were used as housekeeping genes.

#### Enzyme-linked immunosorbent assay (ELISA)

To quantify secreted cytokines from stimulated T cells Human IL-13 ELISA Development Kit (Peprotech, Fisher Scientific) was used according to the manufacturer’s instructions. Optical density was measured at 450 nm (Molecular Devices Spectramax 340PC). All samples were analysed in duplicate and quantified against the standard curve.

#### Statistical analysis

All data are presented as mean ± standard error of the mean (SEM) unless stated otherwise. Outliers were identified using the ROUT method (Q=1%) and excluded from analysis when appropriate. Normality of data distribution was assessed with the Shapiro-Wilk test. For comparisons between two groups, unpaired Welch’s t-test or Mann-Whitney U test was used. For paired samples (pre-vs. post-operative), paired t-test or Wilcoxon matched-pairs signed rank test was applied. For comparisons of stimulated vs. unstimulated conditions within the same donor, paired tests were used. All statistical analyses were performed using Prism version 9.0 (GraphPad Software, San Diego, CA). P-values < 0.05 were considered statistically significant (*, p<0.05; **, p<0.01; ***, p<0.001; ****, p<0.0001).

## Acknowledgements

We thank all patients and volunteers for their participation, the team of the Adipositaszentrum Erlangen PD Dr. Christian Krautz for providing bariatric samples, Doris Wansch and Frederieke Eschenbacher for all their work from recruiting patients to blood sample and skin swab collection. We thank C. Bogdan for continuous support and reagents. Furthermore, we thank D. Vöhringer and U. Schleicher for reagents. Jochen Mattner and Stefan Wirtz for their scientific input.

## Author contributions

J.G. designed, performed and analysed experiments. A.D. and M.W. contributed to specific experiments.

I.H. co-supervised and analysed experiments, wrote and revised the manuscript. C.S. conceptualized the study; designed, supervised and analysed experiments; and wrote and revised the manuscript.

