## Supplemental data for "Preserved Type 2 Immune Cell Plasticity in Human Obesity and Differential Immune Reconstitution After Bariatric Surgery"

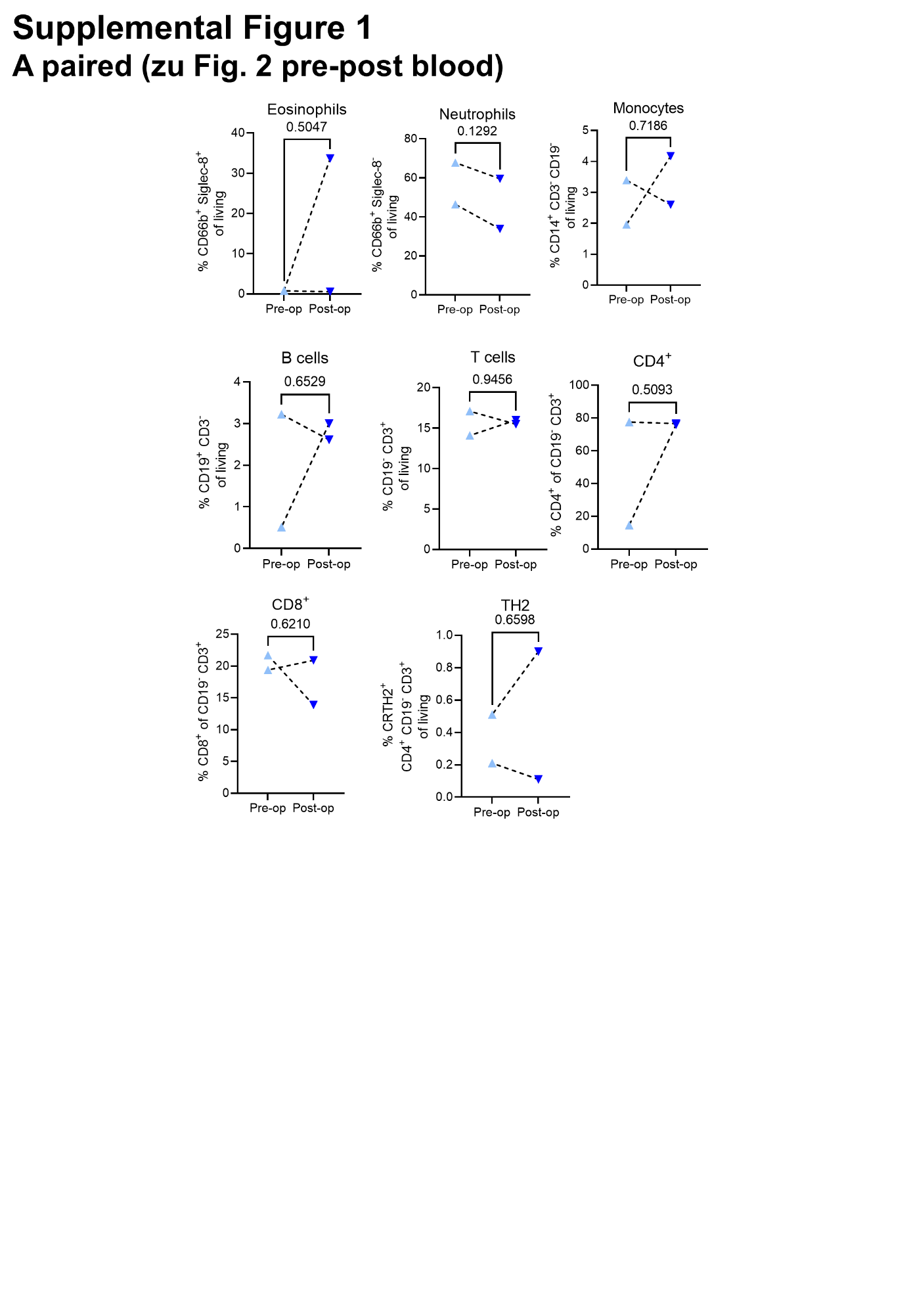

**Supplemental Figure 1.** Paired analysis of blood samples obtained from 2 PWO patients before and after undergoing bariatric surgery. Flow cytometric analysis of blood immune cells from anti-coagulated blood after depletion of erythrocytes with dextran and ACK lysis.

**Supplemental Table 1. Demographic and clinical characteristics of study participants.** Continuous variables are reported as mean ± SD. PWO, people with obesity; BMI, body mass index; GORD, gastro-oesophageal reflux disease.

|  | **PWO (n = 25)** | **Control (n = 29)** | **Lean surgical (n = 8)** |
| --- | --- | --- | --- |
| Sex (male/female) | 15/10 | 14/15 | 5/3 |
| Age (years) | 48.0 ± 8.9 | 30.6 ± 5.9 | 57 ± 20 |
| Weight (kg) | 150.7 ± 29.3 | 72.1 ± 11.6 | 69.2 ± 9.0 |
| BMI (kg/m²) | 48.3 ± 6.9 | 23.5 ± 2.6 | 24.3 ± 3.1 |
| **Surgery type, n (%)** |  |  |  |
| Sleeve gastrectomy | 14 (56) | - | - |
| Gastric bypass | 8 (32) | - | - |
| Revision | 2 (8) | - | - |
| Balloon extraction | 1 (4) | - | - |
| Reflux surgery | - | - | 6 (75) |
| Gastric pacemaker | - | - | 2 (25) |
| **Comorbidities, n (%)** |  |  |  |
| Arterial hypertension | 15 (60) | - | - |
| Type 2 diabetes mellitus | 10 (40) | - | - |
| Mental disorders | 10 (40) | - | - |
| Metabolic disorders | 9 (36) | - | - |
| Sleep apnoea | 13 (52) | - | - |
| GORD | 12 (48) | - | - |
| Respiratory diseases | 8 (32) | - | - |
| Orthopaedic diseases | 17 (68) | - | - |
| Obesity since childhood | 24 (96) | - | - |
| Genetic predisposition | 19 (76) | - | - |
|  | **Pre-OP (n = 13)** | **Post-OP (n = 19)** | **Mean change** |
| Weight (kg) | 171.3 ± 34.6 | 132.3 ± 22.5 | - 32.0 ± 14.1 |
| BMI (kg/m²) | 54.3 ± 6.0 | 43.6 ± 5.6 | - 10.2 ± 4.6 |

**Supplemental Table 2: Antibodies used for flow cytometry.**

| **Antigen** | **Antibody** | **Clone** | **Source** |
| --- | --- | --- | --- |
| *Blood immune cells/ WAT panel* | | | |
| CD3 | BUV737 | OKT3 | Invitrogen |
| CD4 | PerCP-Cy5.5 | OKT4 | BioLegend |
| CD8 | Brilliant Violet 785 | SK1 | BioLegend |
| CD14 | BUV395 | MOP9 | BD Biosciences |
| CD19 | BV421 | HIB19 | BioLegend |
| CD66b | PE/Cy7 | G10F5 | BioLegend |
| CD127 (IL-7Ra) | APC/Fire750 | A019D5 | BioLegend |
| CD161 | APC | HP-3G10 | BioLegend |
| CD294 (CRTH2) | PE/Dazzle 594 | BM16 | BioLegend |
| Siglec-8 | PE | 7C9 | BioLegend |
| Lineage Cocktail 1: |  |  |  |
| CD3 | FITC | SK7 | BD Biosciences |
| CD14 | FITC | MOP9 | BD Biosciences |
| CD16 | FITC | 3G8 | BD Biosciences |
| CD19 | FITC | SJ25C1 | BD Biosciences |
| CD20 | FITC | L27 | BD Biosciences |
| CD56 | FITC | NCAM16.2 | BD Biosciences |
| *T cell panel* | | | |
| CD4 | PerCP-Cy5.5 | OKT4 | BioLegend |
| CD3 | APC/Fire 750 | OKT3 | BioLegend |
| CD25 | BV711 | BC96 | BioLegend |
| PD1 | PE/Cy7 | EH12.2H7 | BioLegend |
| Gata3 | PE | TWAJ | Thermo Scientific |
| FoxP3 | PE/Dazzle 594 | 206D | BioLegend |
| Tbet | APC | 4B10 | BioLegend |
| RorγT | Alexa Fluor 488, FITC | Q21-559 | BD Biosciences |
| *Macrophage panel* | | | |
| CD16 | PE/Dazzle 594 | 3G8 | BioLegend |
| CD64 | FITC | S18012C | BioLegend |
| CD206 | BV786 | 15-2 | BioLegend |
| CD274 (PD-L1) | PE/Cy7 | 29E.2A3 | BioLegend |
| *Eosinophil panel* | | | |
| CD16 | PE/Dazzle 594 | 3G8 | BioLegend |
| CD66b | PE/Cy7 | G10F5 | BioLegend |
| Siglec-8 | PE | 7C9 | BioLegend |

**Supplemental Table 3: Primers used for RT-qPCR.**

| **Target** | **Sequence 5’ 🡪 3’** |
| --- | --- |
| *18S rRNA* | gta acc cgt tga acc cca tt |
| *RPL13A* | gtt ggt gtt cat ccg ctt gc |
| *RETN* | ctg ttg gtg tct agc aag acc |
| *LEP* | tgc ctt cca gaa acg tga tcc |
| *ADIPOQ* | aac atg ccc att cgc ttt acc |
| *PPARG* | tac tgt cgg ttt cag aaa tgc c |
| *IL13* | cct cat ggc gct ttt gtt gac |
| *TNF* | cct ctc tct aat cag ccc tct g |
| *IFNG* | tcg gta act gac ttg aat gtc ca |
| *TBET* | ctg gat gcg cca gga agt tt |
| *GATA3* | gcc cct cat taa gcc caa g |
| *CCR3* | tgg cat gtg taa gct cct ctc |
| *CD44* | ctg ccg ctt tgc agg tgt a |
| *CSF2* | agc ggc ttc agg act ctt g |
| *IRF1* | ctg tgc gag tgt acc gga tg |
| *CD206* | ggg ttg cta tca ctc tct atg c |
| *CD200R* | gag caa tgg cac agt gac tgt t |
